# Genomic heterogeneity affects the response to Daylight Saving Time

**DOI:** 10.1101/2021.03.10.434637

**Authors:** Jonathan Tyler, Yu Fang, Cathy Goldstein, Daniel Forger, Srijan Sen, Margit Burmeister

## Abstract

Circadian rhythms drive the timing of many physiological events in the 24-hour day. When individuals undergo an abrupt external shift (e.g., change in work schedule or travel across multiple time zones), circadian rhythms become misaligned with the new time and may take several days to adjust. Chronic circadian misalignment, e.g., as a result of shift work, has been shown to lead to several physical and mental health problems. Despite the serious health implications of circadian misalignment, relatively little is known about how genetic variation affects an individual’s ability to shift to abrupt external changes. Accordingly, we use the one-hour advance from the onset of daylight saving time (DST) as a natural experiment to comprehensively study how individual heterogeneity affects the shift of sleep-wake rhythms in response to an abrupt external time change. We find that individuals genetically predisposed to a morning tendency adjust to the advance in a few days, while genetically predisposed evening-inclined individuals have not shifted. Observing differential effects by genetic disposition after a one-hour advance underscores the importance of heterogeneity in adaptation to external schedule shifts, and these genetic differences may affect how individuals adjust to jet lag or shift work as well.

## INTRODUCTION

Circadian (about a day) rhythms are fundamental processes in most organisms that time many molecular, physiological, and behavioral events across 24 hours^1^, such as regulating hormone release^2^, body temperature^3^, metabolism^4^, and sleep-wake patterns^5^. In humans, these rhythms are synchronized (i.e., *entrained*) to the environment primarily through the external light-dark cycle. Although the rhythms are aligned to the environment, a key characteristic is that they are free-running and will persist in constant conditions^6^. Thus, circadian rhythms will not instantaneously adjust to abrupt external shifts, e.g., changes in work schedule or travel across multiple time zones.

Notably, an individual’s inherent capacity for realignment of the endogenous circadian rhythm to the external schedule after an abrupt shift has implications in overall individual well-being as well as mental and physical health^7^. For example, temporary and sudden shifts in external time due to traveling or daylight saving time (DST) result in misalignment between the circadian timekeeping system and the external environment^8^. In addition to the misalignment between the central clock and external light-dark schedule and behaviors such as sleeping and eating, desynchronization among peripheral clocks also occurs. Temporary desynchronization leads to impaired sleep at night and excessive daytime sleepiness as well as other consequences, such as increased automotive accidents^9–11^ and heart attacks^12,13^. Importantly, the magnitude of disruption to central and peripheral rhythms that results from abrupt external schedule changes demonstrates significant interindividual variability^14–18^. Therefore, a deeper understanding of interindividual differences in the response to these temporary shifts can help address the broader issue of chronic circadian misalignment.

Previous studies have used the DST transition as a unique opportunity to investigate how individuals adjust to any external time change^19–24^. However, these analyses relied on subjective self-report, limiting the traits that could be captured and often misrepresenting personal sleep information^25^. Moreover, previous studies were limited in power due to small sample sizes and often reported contradictory results^24^. These contradictory results regarding the impact of DST may be a result of studies failing to account for circadian tendency and interindividual heterogeneity in sleep behavior because the capacity to measure sleep objectively over several nights has only recently become feasible.

Now, however, genome-wide association studies provide objective measures that describe predisposition to certain diseases and phenotypes given individual genetic variation^26^. In particular, recent work has revealed genetic associations with sleep midpoint from accelerometer-derived estimates of sleep rather than biased self-reported sleep information^25^. This validated and objective genetic sleep midpoint association provides a surrogate measure to characterize an individual’s genomic predisposition to morningness or eveningness, specifically as it relates to the sleep-wake cycle.

Here, we use the one-hour advance from the onset of daylight saving time as a natural experiment to comprehensively study how individual heterogeneity affects how sleep-wake rhythms shift in response to an abrupt external time change. In total, from a cohort of 831 medical interns^27^ with genotype information, we analyze thousands of sleep events the week before and the week after DST onset in 2019. After analysis, we see that morning-inclined individuals adjust more rapidly to the time shift better than evening-inclined. Moreover, we find that social jet lag^28^ is exaggerated in evening-inclined individuals the week after DST. Ultimately, our results provide a more comprehensive evaluation of how genetic predispositions affect how the sleep-wake rhythm patterns shift in response to an external change. Since DST is only a one-hour advance, observing differential effects by genetic disposition underscores the importance of heterogeneity in more extreme cases such as shift work. Additionally, our results further establish the utility of the polygenic score as a surrogate measure of phenotypes.

Finally, understanding how subtle phenotypic differences affect shifting patterns will help inform individuals on how to prepare for abrupt external shifts (e.g., DST, changes in schedule, transmeridian travel, etc.), and ultimately, continue to inform our understanding of circadian misalignment.

## RESULTS

### Sleep midpoint in the morning group is significantly earlier than that in the evening group

We used the Objective Sleep Midpoint polygenic score (OSM PGS), calculated using a genome-wide chronotype genetic analysis of 85,670 individuals^25^, as an objective measure for chronotype rather than subjective self report. Specifically, we calculated the OSM PGS with the methods described previously^29,30^ for all subjects with genetic information (N=831). As a higher OSM PGS corresponds to a later sleep midpoint, we divided all PGS scores into three quantiles and characterized the top quantile as an evening group and the bottom quantile as a morning group. In Figure 1, using all the sleep events up to the onset of DST, we plot the distribution of sleep midpoints for the two groups on work (e.g., Sunday-Thursday) and free (e.g., Friday-Saturday) nights. As expected, the morning group had significantly earlier sleep midpoints on both free and work nights, and the evening group displayed later sleep midpoints (Figure 1).

**Figure 1:**
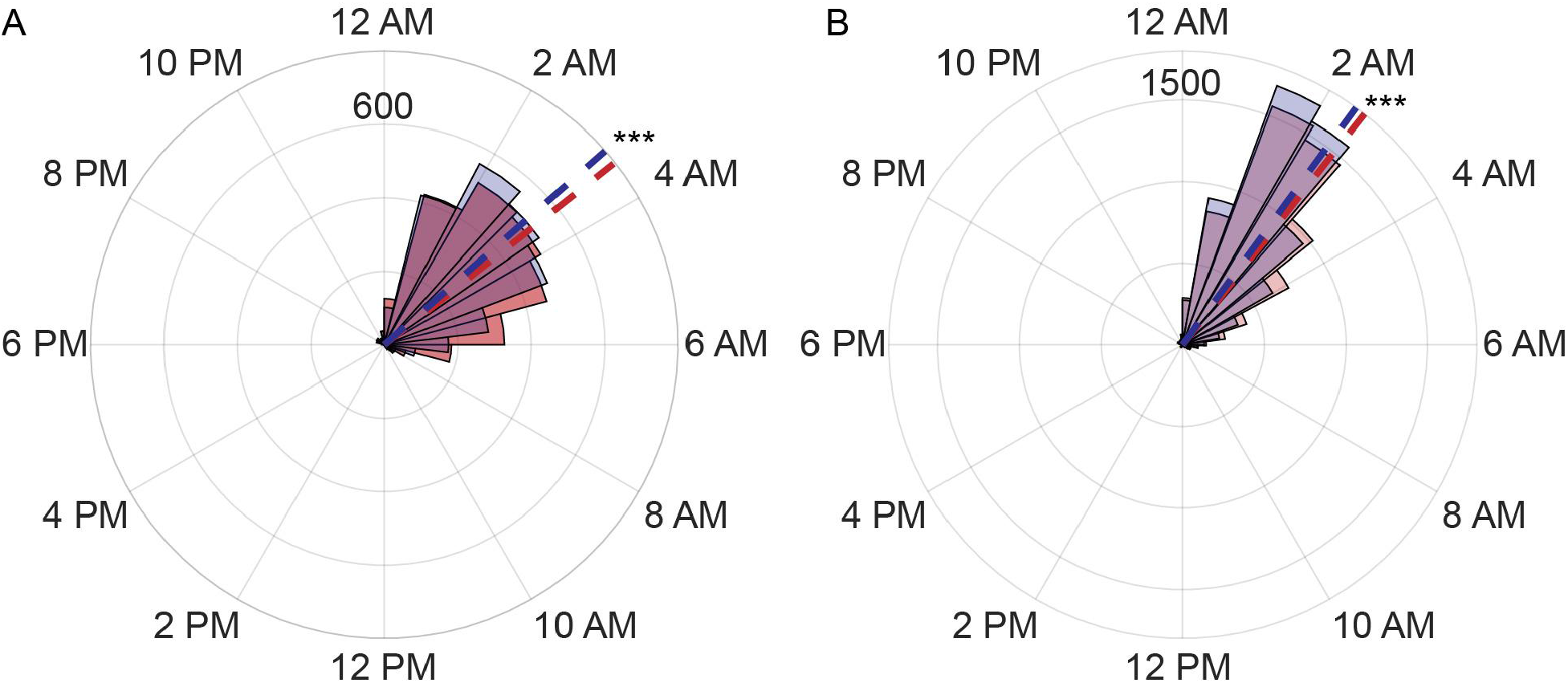
Sleep midpoints are significantly different in the two groups. The distribution of sleep midpoints in the morning (blue shade) group is earlier than that of the evening (red) group on free nights (A) and work nights (B). The mean sleep midpoints are plotted as dashed lines. On free nights (A), the mean sleep midpoint of the morning group is 3:15 AM (n = 2771 sleep events), and the mean sleep midpoint of the evening group is 3:26 AM (n = 2754 sleep events, P = 0.0003). On work nights (B), the mean sleep midpoint of the morning group is 2:25 AM (n = 7296), and the mean sleep midpoint of the evening group is 2:33 AM (n = 7227 sleep events, P = 7.2e-6).

### Evening population sleeps less the week following DST relative to the week before

For the week before and after DST onset, we computed the mean time asleep, in minutes, for both the morning and evening groups (Figure 2A). We saw a slight increase in the average time asleep in the morning population (mean 362 +/- 131 before and 367 +/- 128 minutes after DST, Figure 2A). Thus, the morning population, in general, was able to shift their sleep-wake schedule to align with the one-hour shift forward. In contrast, the mean time asleep decreased in the evening population (from 367 +/- 128 minutes before to 359 +/- 132 minutes the week after, Figure 2A), close to a level of significance (P = 0.16). Altogether, the group with the later circadian tendency could not successfully shift their sleep-wake schedule to align with the one-hour shift forward.

**Figure 2:**
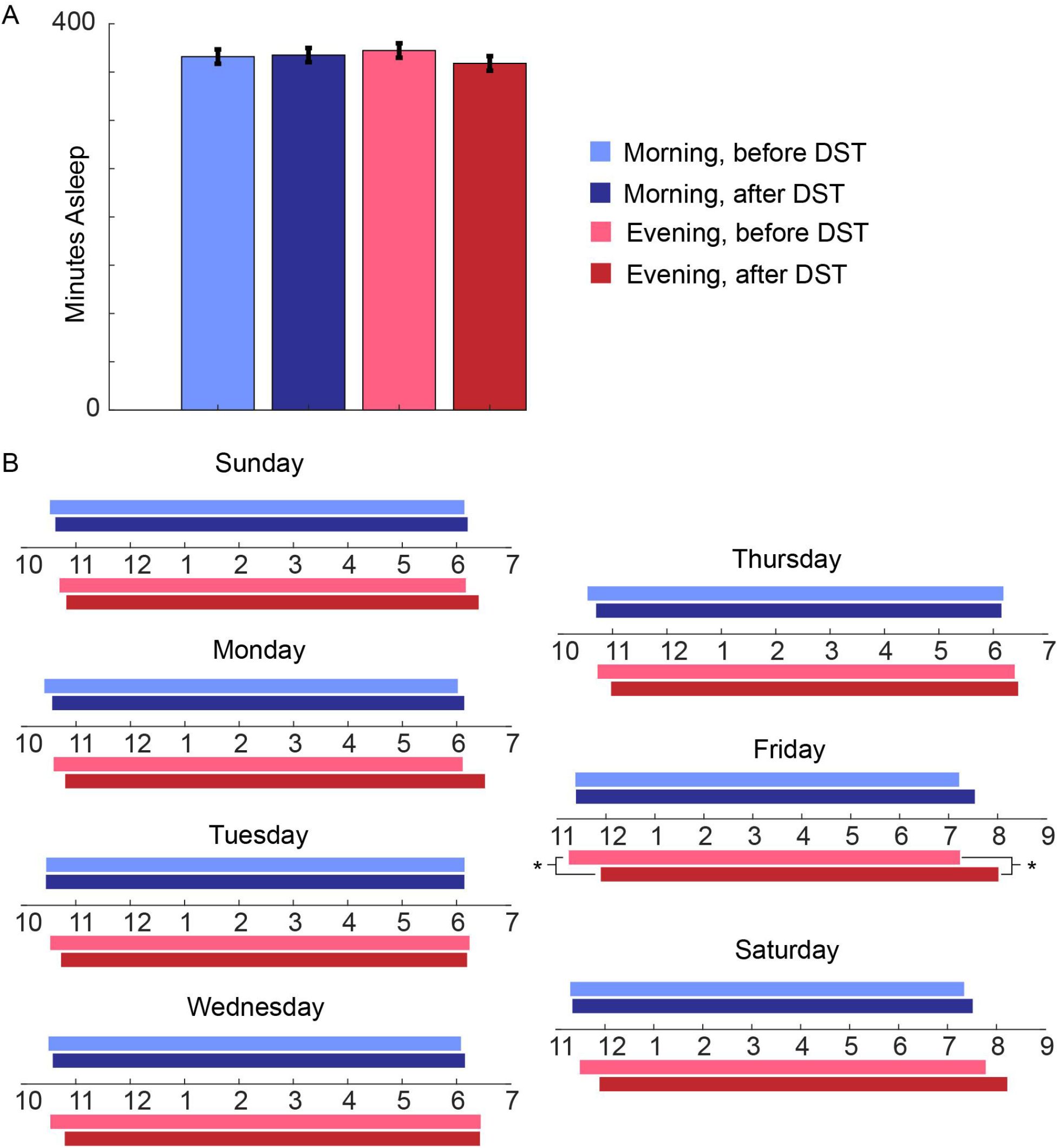
Analysis of Sleep Events before and after DST starts. (A) The mean asleep time for the morning population is consistent across the week before DST (n = 1108 sleep events) and the week after DST (n = 1216 sleep events). The mean asleep time for the evening population decreases (n = 1175 sleep events) relative to the week before DST (n = 1021 sleep events). (B) Mean sleep profiles of the morning (blue) and evening (red) populations for the night of the week before (lighter shade) and after DST (darker shade). For statistical analysis on circular data, we use the Parametric Watson-Williams multisample test for equal means (P = 0.01999 for sleep onset and 0.0161 for sleep offset). See Table 1 for the sample sizes in each group.

### Morning population shifts to the new time in three days while the evening population has not shifted one week after DST

For both populations, we computed the mean sleep onset and offset times for each day of the week before and after DST (Figure 2B). In the morning population, sleep onset times are delayed on the Sunday and Monday nights after DST (Figure 2B). However, after Monday night, the sleep profiles of the morning population are nearly identical across the week before and after DST. Moreover, on the more free nights (e.g., Friday and Saturday), the morning population sleep profiles continue to remain nearly identical across the two weeks.

The evening type population also exhibited sleep onset times that are delayed on the Sunday and Monday nights after DST. Furthermore, unlike the morning population, the evening population continued to exhibit later sleep onset and offset times the week after DST compared with the week before DST (Figure 2B). Importantly, the sleep onset times display a more dramatic delay relative to the week before than the offset times, which are likely fixed due to work schedule constraints. Because work schedule constraints are less on weekends, the differences in sleep profiles are more dramatic on the Friday and Saturday night after DST. In fact, on Friday, there is a significant difference in the sleep onset time of around 45 minutes and in the sleep offset time of around 40 minutes (Figure 2B). Therefore, even one week after DST, the evening population has not shifted to the new external time.

### Social jet lag significantly increases in the evening population the week after DST

Finally, we assessed social jet lag, the difference between the sleep midpoint on workdays and free days^28,31^, the weeks before and after DST in each population (Figure 3A). Specifically, we grouped the sleep midpoints of Monday-Thursday nights before and after DST as the work-night sleep midpoints. Similarly, we group the sleep midpoints on Friday and Saturday nights as the free-night sleep midpoints.

**Figure 3:**
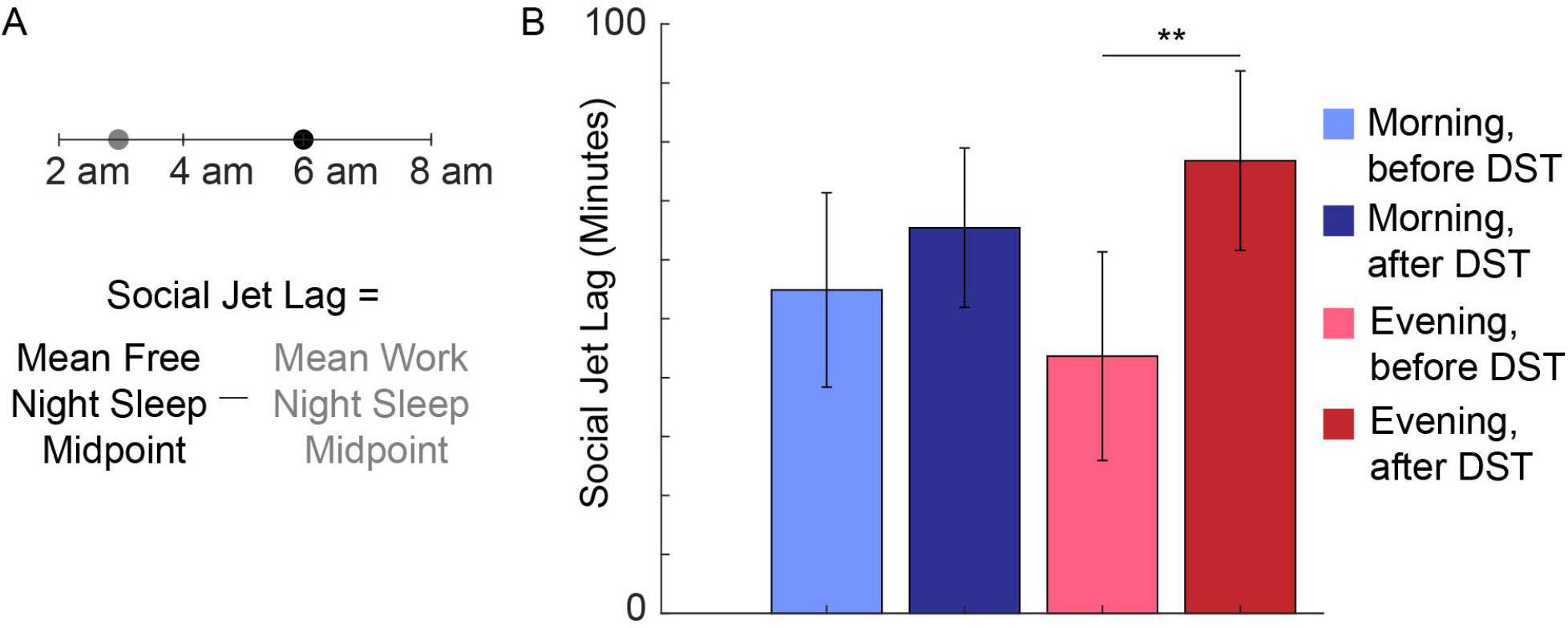
Social jet lag is significantly higher in evening population after DST. (A) Schematic of the social jet lag calculation. For each individual with at least one work night sleep event and one free night sleep event, we calculate the mean sleep midpoint for the workweek and the mean sleep midpoint for the free nights. Then, social jet lag is defined as the difference between the mean free night sleep midpoint and the mean work night sleep midpoint. (B) For the morning population, social jet lag is consistent across the week before DST (n = 133 subjects) and the week after DST (n = 141 subjects). For the evening population, social jet lag significantly decreased (P = 0.0051) the week after DST (n = 134 subjects) compared with the week before DST (n = 118 subjects).

As expected, we see that social jet lag is significantly increased in the evening population (from 44 to 77 minutes, Figure 3B) after DST. Thus, even though the evening population continues to have a later sleep onset each night of the week following DST (Figure 2B), the difference in midpoint from the work nights to the following Friday is significantly more pronounced. In contrast, we see only a small increase in the social jet lag of the morning population (55 to 65 minutes, Figure 3B) after DST, providing further evidence that the morning population has successfully shifted to time change the weekend following DST.

## DISCUSSION

Here, we performed a comprehensive analysis of how individual heterogeneity affects an individual’s ability to adjust to an abrupt shift. We used the Objective Sleep Midpoint polygenic score^25,29,30^ to characterize individual heterogeneity in sleep-wake rhythms. In particular, we labeled morning-inclined individuals as those with an OSM PGS in the bottom third of all scores, and we labeled evening-inclined individuals as those with an OSM PGS in the top third of all scores. This characterization revealed significant differences in sleep midpoints on both free and work nights, similar in magnitude to other reports^25^. Thus, we used the OSM PGS as a surrogate marker for chronotype.

In summary, we saw dramatic differences in adaptation to the one hour DST time shift between individuals genetically predisposed to a morning versus evening circadian inclination. First, the evening group exhibited lower average time asleep in the week following DST relative to the week before, corroborating previous work which reported a decrease in sleep duration the week after DST relative to the previous week^22^. The lower average time asleep is particularly concerning in a population of medical interns, for which it has previously been shown that they are already sleep-deprived^32^. Additional lack of sleep the week after DST in the evening group may exaggerate depressive symptoms, more likely in evening individuals^33^, or increase medical errors.

Further analysis revealed that both groups exhibited the expected delay in sleep onset and offset for the Sunday and Monday nights after DST onset. Thus, even though there may be a build-up of sleep pressure due to less sleep the Saturday night of DST onset, contrary to a previous report^21^, it is not enough to result in an instantaneous shift. After Monday, the sleep timing of the morning population realigned to the sleep schedule before DST. This indicates that the shift of the morning population is not a masking effect of the workweek, but rather an actual shift to the new time in both the sleep-wake rhythm and inherent biological clock. Morning individuals may shift so quickly because these individuals have a more rapid build-up of sleep loss, and thus, after an advance, they can shift their bedtimes earlier^34^.

In contrast, the evening population sleep profiles continued to exhibit a delay in sleep onset for the rest of the workweek. Furthermore, on free nights a week after DST (i.e., Friday and Saturday), the evening population showed a drastic delay in both sleep onset and offset relative to the week before DST. Thus, the workweek schedule masks the magnitude to which the sleep-wake rhythms in the evening population are misaligned to the new time. Assessment of sleep-wake timing on free days demonstrated that the evening population has not shifted to the time change one week after DST onset. Moreover, given fixed wake-up times due to the interns’ rigid work requirements, the evening population had a significant increase in social jet lag the week after DST onset relative to the week before. This result further supports the difficulty evening individuals have in phase advancing to realign with external time in response to the DST time shift.

The significant increase in social jet lag highlights the desynchronization between the independent homeostatic sleep-wake and internal circadian sleep rhythms in the evening population. The sleep-wake rhythm drive, as seen in the work nights, does not display as dramatic of a shift because of the increase in sleep pressure. However, the internal biological clock drive, seen in the phase of the free nights, continues to be delayed one week after DST in the evening population. As the desynchronization between the two rhythms remains relatively constant in the morning population, the more extreme desynchronization in the two sleep rhythms in the evening population underscores that the magnitude of desynchronization between peripheral and central clocks displays significant interindividual variability. The evening population’s decreased ability to mitigate abrupt shifts may result in a higher likelihood of adverse health events or medical errors.

We acknowledge that our study is limited in that all participants are from a cohort of medical Interns. While this limits generalizability, it also provides the advantage that the age range is limited, avoiding having to account for age effects on circadian behavior. Another limitation is that we do not consider the geographical location of the subjects. Dawn times vary widely around the spring equinox, and thus, around the spring DST transition with a differential effect across latitudes^24^. Therefore, individuals at certain latitudes may mitigate the shift better than at other latitudes because of stronger sunlight at earlier dawn times.

Altogether, our results demonstrate the power of using polygenic scores in differentiating individual genomic predisposition to morningness or eveningness and how these characterizations reveal interindividual variability in shifting to external changes. Observing significant differential effects after a one-hour external shift emphasizes that genetic variation may play an important role in mitigating more extreme shits, and thus, how individuals respond to chronic misalignment due to shift work and the extent to which individuals suffer mental and physical health problems as a result.

## METHODS

### Participants

The Intern Health Study (IHS) is a multi site cohort study that follows physicians across several institutions in their first year of residency. Interns entering residency in 2018 The intern participants entered residency in 2018 and were contacted via email 2-3 months before residency onset regarding details of the study and provided informed written consent to participate. Interns that consented were given a $25 gift certificate upon completion of a baseline survey and another $25 gift certificate after completing a follow-up survey ^27,35^. All subjects were invited to wear Fitbits to track sleep patterns, heart rate, and physical activity. Both the IRB at the University of Michigan and participating institutions approved the study, and all work was carried out in accordance with these approvals and all relevant regulations.

### Sleep event processing

We characterized each sleep event as a nighttime sleep event if it started between the hours of 6 pm and 6 am the following morning. Otherwise, the sleep event was characterized as a daytime sleep event. Furthermore, we classified each sleep event by the day of the week on which it started. For example, a sleep event that started after 6 pm on Sunday but before 6 am the following Monday was classified as a Sunday nighttime sleep event even if it technically started on Monday.

To assess the ability to shift to the external time change, we further refined the data set, following the convention introduced in previous studies examining sleep changes after DST^19,21^. In particular, we characterized sleep events as being the week before DST if they were nighttime sleep events from Saturday, March 2nd, to Friday, March 8th. Similarly, we characterized sleep events as being the week after DST onset if they were nighttime sleep events from Sunday, March 10th, to Saturday, March 16th. In this way, we guaranteed, as much as possible, a uniform representation of sleep events starting each night of the week. Table 1 reports a breakdown of the number of sleep events each night of the week both before and after DST onset.

**Table 1:**
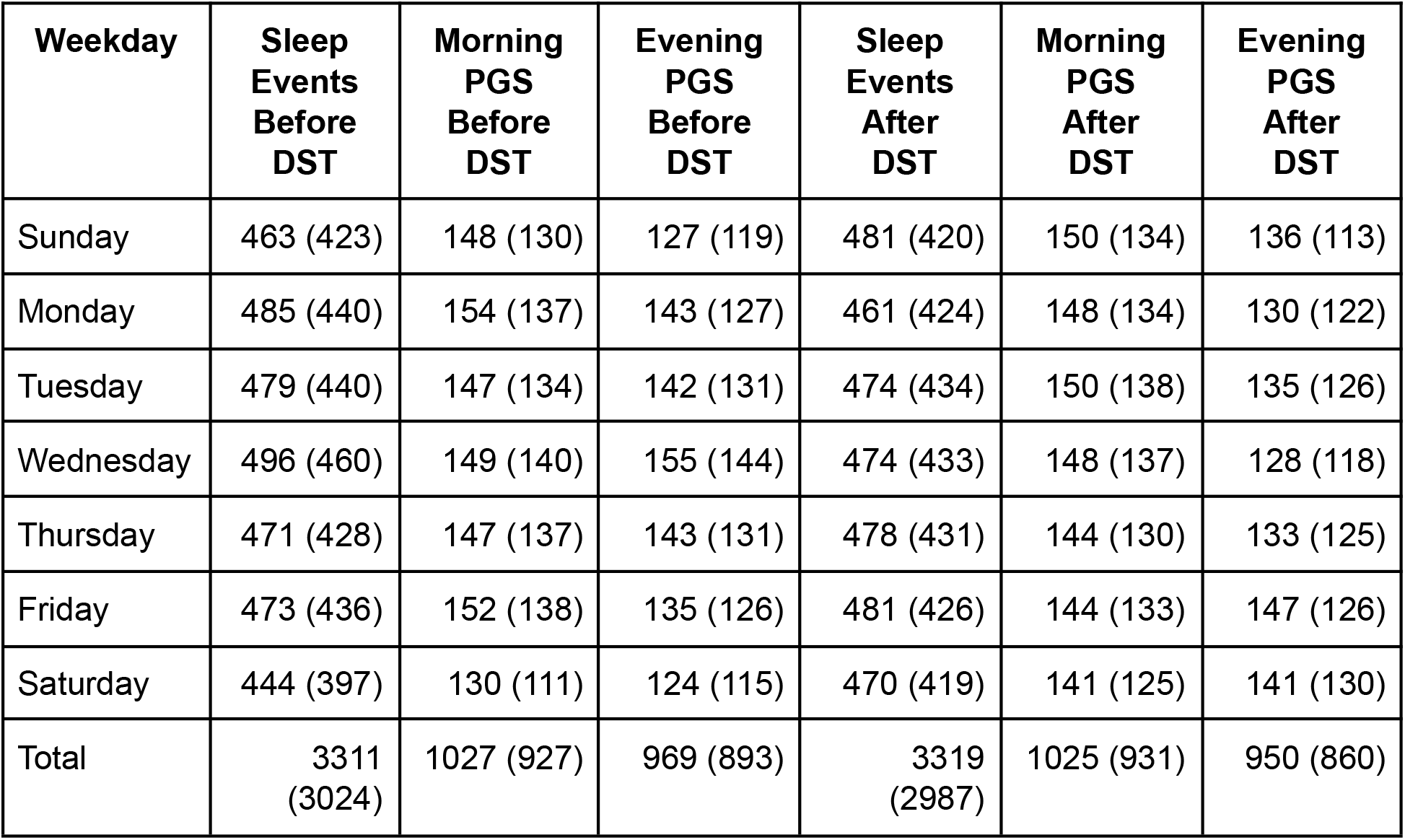
Number of weekday sleep events before and after DST. Here, we list the total number of nighttime sleep events the weeks before and after DST onset starting on the respective day of the week (columns 2 and 5). Furthermore, we list the number of sleep events of the morning population (before DST, column 3 and after DST, column 6) and the evening population (columns 4 and 7). The values listed in parentheses are the number of sleep events that are greater than four hours.

We then subdivided the sleep events both before and after DST onset based on whether the subject was labeled as morning- or evening-inclined. In Table 1, we breakdown the number of sleep events in the morning and evening groups for the weeks before and after DST. In the end, we analyzed 1027 (969) sleep events the week before DST and 1025 (950) sleep events the week after DST for the morning and evening groups (Table 1). These sleep events were distributed uniformly across the seven week nights in each population (Table 1).

To account for the effect of shift schedules, we only used sleep events from subjects that had sleep events in both weeks on the same day of the week. I.e., for the Monday night sleep analysis, we only took sleep events from subjects that had a sleep event the Monday before and the Monday after DST. Furthermore, we rejected any sleep event that was less than four hours long to remove any bias due to interruptions in a normal sleep event (e.g., an interruption because the subject was called into work in the middle of the night). In this way, we removed, as much as possible, any bias introduced into the data from shift changes. Altogether, after removing sleep events that were less than four hours, we lost about ten percent of the sleep events. See the parentheses values in Table 1 for the number of sleep events that were longer than four hours, and thus, the total number of sleep events used in the analysis

Finally, at least one work night sleep event and one free night sleep event were required for inclusion of individuals in the social jet lag analysis.

### Genotyping and PGS calculation

We collected DNA from 2018 cohort subjects (n = 1,624) using DNA Genotek Oragene Mailable Tube (OGR-500)^36^ through the mail. DNA was extracted and genotyped on Illumina Infinium CoreExome-24 with v.1.3 Chips, containing 595,427 SNPs. We then implemented a quality check of genotype data with PLINK v.1.9 (www.cog-genomics.org/plink/1.9/)^37^. Samples with call rate < 99% (n = 18) or with a sex mismatch between genotype data and reported data (n = 4) were excluded. Duplicated samples (n = 3) with lower call rate was excluded. We only included SNPs on autosomal chromosomes, with call rate ≥ 98% (after sample removal) and minor allele frequency (MAF) ≥ 0.005. A total of 331,077 SNPs and 1,599 samples were considered for further analysis. Of the 1,599 samples, we analyzed 831 that had corresponding wearable data. The samples contained mixed ancestries to increase sample size and power.

To calculate the PGS of objective sleep midpoint, we used the GWAS summary statistics of accelerometer-based sleep midpoint from UK Biobank containing 85,670 samples^25^. We used PRSice v.2^29^ to calculate OSM-PGS for our intern subjects. The OSM-PGS was calculated using all the 225,466 variants genotyped in our sample (with MAF ≥ 0.1 and outside the major histocompatibility complex (MHC) region) that overlap with summary statistic data from the OSM GWAS (i.e., p-value thresholds = 1). The OSM-PGS was then calculated with a weighted additive model:

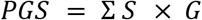

where S is the MDD GWAS summary statistic effect size for the effect allele and G = 0,1,2 is the number of effect alleles observed. The OSM-PGS was then mean-centred and scaled to 1 SD.

### Data and statistical analysis

All data and statistical analyses were performed in Matlab R2019b. To analyze the mean time asleep in the week before and after DST, we aggregated all sleep profiles from the 831 subjects with PGS and wearable data in the respective weeks. We then computed the mean time asleep for the week before DST by averaging the time asleep for all nighttime sleep events from Saturday, March 2, 2019 to Friday, March 8, 2019. We performed the same calculation for the average time asleep the week after DST (nighttime sleep events from

Sunday, March 10, 2019 to Saturday, March 16, 2019). Finally, we performed a two-sample t-test at a significance level of 95% to assess whether mean time asleep was significantly different before and after DST in the two groups independently.

We accounted for the circular nature of sleep data by using circular statistic functions from the Circular Statistics Toolbox in Matlab^38^. For example, we saved all sleep start times and normalized them to a value between 0 and 2π (0 corresponds to midnight and π corresponds to noon) and computed the mean sleep midpoints using the circular mean function. Then, we assessed significance by performing a Parametric Watson-Williams multi-sample significance test for unequal means at a significance level of 95%. In a similar way, we computed the mean sleep onset and sleep end times for both the morning and evening groups before and after DST and assessed significance in each case.

To analyze social jet lag, we first found all subjects that had at least one work night (Monday-Thursday) sleep event and one free night (Friday-Saturday) sleep event the week before DST. We then calculated the social jet lag for the individual as the mean sleep midpoint on free nights minus the mean sleep midpoint on work nights. We performed the same procedure for the week after DST. Note that we did not require subjects to be in both the week before and the week after, just one or the other. Then, we computed the mean social jet lag for each group before and after DST and assessed significance using a two-sample t-test at a significance level of 95%.

## Acknowledgements

We would like to thank the medical interns that participated in this study. JT is grateful for support from a National Institutes of Health Training Grant (T32 HL007622). The Intern Health Study and genotyping are supported by the National Institute of Mental Health, MH101459.

## Data availability

The de-identified data from Intern Health Study is available through the Psychiatric Genomics Consortium (PGC):https://www.med.unc.edu/pgc/shared-methods. PGC phase 2–UK Biobank–23andMe MDD GWAS meta-analysis summary statistics: https://www.nature.com/articles/s41593-018-0326-7#data-availability.

## REFERENCES

1. Dunlap, J. C. Molecular bases for circadian clocks. Cell 96, 271–290 (1999).

2. Gnocchi, D. & Bruscalupi, G. Circadian Rhythms and Hormonal Homeostasis: Pathophysiological Implications. Biology 6, (2017).

3. Aschoff, J. Circadian control of body temperature. J. Therm. Biol. 8, 143–147 (1983).

4. Fonken, L. K. et al. Light at night increases body mass by shifting the time of food intake. Proc. Natl. Acad. Sci. U. S. A. 107, 18664–18669 (2010).

5. Czeisler, C. A., Weitzman, E. d., Moore-Ede, M. C., Zimmerman, J. C. & Knauer, R. S. Human sleep: its duration and organization depend on its circadian phase. Science 210, 1264–1267 (1980).

6. Takahashi, J. S., Hong, H.-K., Ko, C. H. & McDearmon, E. L. The genetics of mammalian circadian order and disorder: implications for physiology and disease. Nat. Rev. Genet. 9, 764–775 (2008).

7. Roenneberg, T. & Merrow, M. The Circadian Clock and Human Health. Curr. Biol. 26, R432–43 (2016).

8. Vosko, A. M., Colwell, C. S. & Avidan, A. Y. Jet lag syndrome: circadian organization, pathophysiology, and management strategies. Nat. Sci. Sleep 2, 187 (2010).

9. Ferguson, S. A., Preusser, D. F., Lund, A. K., Zador, P. L. & Ulmer, R. G. Daylight saving time and motor vehicle crashes: the reduction in pedestrian and vehicle occupant fatalities. Am. J. Public Health 85, 92–95 (1995).

10. Lambe, M. & Cummings, P. The shift to and from daylight savings time and motor vehicle crashes. Accid. Anal. Prev. 32, 609–611 (2000).

11. Varughese, J. & Allen, R. P Fatal accidents following changes in daylight savings time: the American experience. Sleep Med. 2, 31–36 (2001).

12. Sandhu, A., Seth, M. & Gurm, H. S. Daylight savings time and myocardial infarction. Open Heart 1, e000019 (2014).

13. Manfredini, R. et al. Daylight Saving Time and Acute Myocardial Infarction: A Meta-Analysis. J. Clin. Med. Res. 8, (2019).

14. Folkard, S. Do permanent night workers show circadian adjustment? A review based on the endogenous melatonin rhythm. Chronobiol. Int. 25, 215–224 (2008).

15. Stone, J. E. et al. Temporal dynamics of circadian phase shifting response to consecutive night shifts in healthcare workers: role of light-dark exposure. J. Physiol. 596, 2381–2395 (2018).

16. Kervezee, L., Cermakian, N. & Boivin, D. B. Individual metabolomic signatures of circadian misalignment during simulated night shifts in humans. PLoS Biol. 17, e3000303 (2019).

17. Koshy, A., Cuesta, M., Boudreau, P., Cermakian, N. & Boivin, D. B. Disruption of central and peripheral circadian clocks in police officers working at night. FASEB J. 33, 6789–6800 (2019).

18. Kervezee, L., Kosmadopoulos, A. & Boivin, D. B. Metabolic and cardiovascular consequences of shift work: The role of circadian disruption and sleep disturbances. Eur. J. Neurosci. 51, 396–412 (2020).

19. Monk, T. H. & Folkard, S. Adjusting to the changes to and from Daylight Saving Time. Nature 261, 688–689 (1976).

20. Nicholson, A. N. & Stone, B. M. Adaptation of sleep to British Summer Time [proceedings]. J. Physiol. 275, 22P–23P (1978).

21. Monk, T. H. & Aplin, L. C. Spring and autumn daylight saving time changes: studies of adjustment in sleep timings, mood, and efficiency. Ergonomics 23, 167–178 (1980).

22. Lahti, T. A., Leppämäki, S., Lönnqvist, J. & Partonen, T. Transition to daylight saving time reduces sleep duration plus sleep efficiency of the deprived sleep. Neurosci. Lett. 406, 174–177 (2006).

23. Lahti, T. A. et al. Transition into daylight saving time influences the fragmentation of the rest-activity cycle. J. Circadian Rhythms 4, 1 (2006).

24. Kantermann, T., Juda, M., Merrow, M. & Roenneberg, T. The human circadian clock’ sseasonal adjustment is disrupted by daylight saving time. Curr. Biol. 17, 1996–2000 (2007).

25. Jones, S. E., van Hees, V. T. & Mazzotti, D. R. Genetic studies of accelerometer-based sleep measures yield new insights into human sleep behaviour. Nature (2019).

26. Tam, V. et al. Benefits and limitations of genome-wide association studies. Nat. Rev. Genet. 20, 467–484 (2019).

27. Sen, S. et al. A prospective cohort study investigating factors associated with depression during medical internship. Arch. Gen. Psychiatry 67, 557–565 (2010).

28. Wittmann, M., Dinich, J., Merrow, M. & Roenneberg, T. Social jetlag: misalignment of biological and social time. Chronobiol. Int. 23, 497–509 (2006).

29. Euesden, J., Lewis, C. M. & O’Reilly, P. F. PRSice: Polygenic Risk Score software. Bioinformatics 31, 1466–1468 (2015).

30. Fang, Y., Scott, L., Song, P., Burmeister, M. & Sen, S. Genomic prediction of depression risk and resilience under stress. Nat Hum Behav 4, 111–118 (2020).

31. Roenneberg, T., Wirz-Justice, A. & Merrow, M. Life between clocks: daily temporal patterns of human chronotypes. J. Biol. Rhythms 18, 80–90 (2003).

32. Kalmbach, D. A. et al. Effects of Sleep, Physical Activity, and Shift Work on Daily Mood: a Prospective Mobile Monitoring Study of Medical Interns. J. Gen. Intern. Med. 33, 914–920 (2018).

33. Kivelä, L., Papadopoulos, M. R. & Antypa, N. Chronotype and Psychiatric Disorders. Curr Sleep Med Rep 4, 94–103 (2018).

34. Dijk, D.-J. & Archer, S. N. PERIOD3, circadian phenotypes, and sleep homeostasis. Sleep Med. Rev. 14, 151–160 (2010).

35. Guille, C. et al. Web-Based Cognitive Behavioral Therapy Intervention for the Prevention of Suicidal Ideation in Medical Interns: A Randomized Clinical Trial. JAMA Psychiatry 72, 1192–1198 (2015).

36. Rogers, N. L., Cole, S. A., Lan, H.-C., Crossa, A. & Demerath, E. W. New saliva DNA collection method compared to buccal cell collection techniques for epidemiological studies. Am. J. Hum. Biol. 19, 319–326 (2007).

37. Chang, C. C. et al. Second-generation PLINK: rising to the challenge of larger and richer datasets. Gigascience 4, 7 (2015).

38. Berens, P. CircStat: A MATLAB Toolbox for Circular Statistics. Journal of Statistical Software, Articles 31, 1–21 (2009).

